# Reduced infectivity but increased immune escape of the new SARS-CoV-2 variant of concern Omicron

**DOI:** 10.1101/2021.12.24.474110

**Authors:** Jie Hu, Pai Peng, Kang Wu, Quan-xin Long, Juan Chen, Kai Wang, Ni Tang, Ai-long Huang

## Abstract

A new detected SARS-CoV-2 variant Omicron (B.1.1.529) had reported from more than 80 countries. In the past few weeks, a new wave of infection driven by Omicron is in progress. Omicron Spike (S) protein pseudotyped virus was used to determine the effect of S mutations on its capacity of infectivity and immune evasion. Our results showed the lower entry efficiency and less cleavage ability of Omicron than D614G variant. Pseudotype-based neutralizing assay was performed to analyze neutralizing antibodies elicited by previously infection or the RBD-based protein subunit vaccine ZF2001 against the Omicron variant. Sera sampled at around one month after symptom onset from 12 convalescents who were previously infected by SARS-CoV-2 original strain shows a more than 20-fold decrease of neutralizing activity against Omicron variant, when compared to D614G variant. Among 12 individuals vaccinated by RBD subunit vaccine, 58.3% (7/12) sera sampled at 15-60 days after 3rd-dose vaccination did not neutralize Omicron. Geometric mean titers (GMTs, 50% inhibitory dose [ID50]) of these sera against Omicron were 9.4-fold lower than against D614G. These results suggested a higher risk of Omicron breakthrough infections and reduced efficiency of the protective immunity elicited by existing vaccines. There are important implications about the modification and optimization of the current epidemic prevention and control including vaccine strategies and therapeutic antibodies against Omicron variant.

## Introduction

On 24 November, a new detected variant B.1.1.529 of SARS-CoV-2 by South Africa was reported to WHO. After only two days, this variant was designated as “Variant of Concern” (VOC) and named as Omicron. In the past few weeks, Omicron had reported from more than 80 countries. It has been reported as the dominant SARS-CoV-2 in U.S. due to the rapid spread of Omicron. A new wave of infection driven by Omicron is in progress.

The major reason that Omicron raise a great concern is its accumulated mutations, including more than 30 of those in the spike (S) protein. More importantly, 15 of those mutations occurs on receptor-binding domain (RBD) (Fig. 1a), which is not only the vital binding site to the host receptor angiotensin-converting enzyme 2 (ACE2) for the entry of SARS-CoV-2, but also the key target of neutralizing antibodies produced by immune response and therapeutics antibodies. By contrast, other VOCs including Alpha, Beta, Gamma and Delta possess 9-12 mutations on their S protein regions. Even so, some crucial mutations like D614G, N501Y, K417N and E484A in these known VOCs have been reported about the effect on viral infectivity and transmission^1,2,3,4^. Except for these shared mutations mentioned above, with additional mutations on Omicron, there is a pressing need for more evidence directed at the evaluation of the synergistic effect.

**Fig.1.**
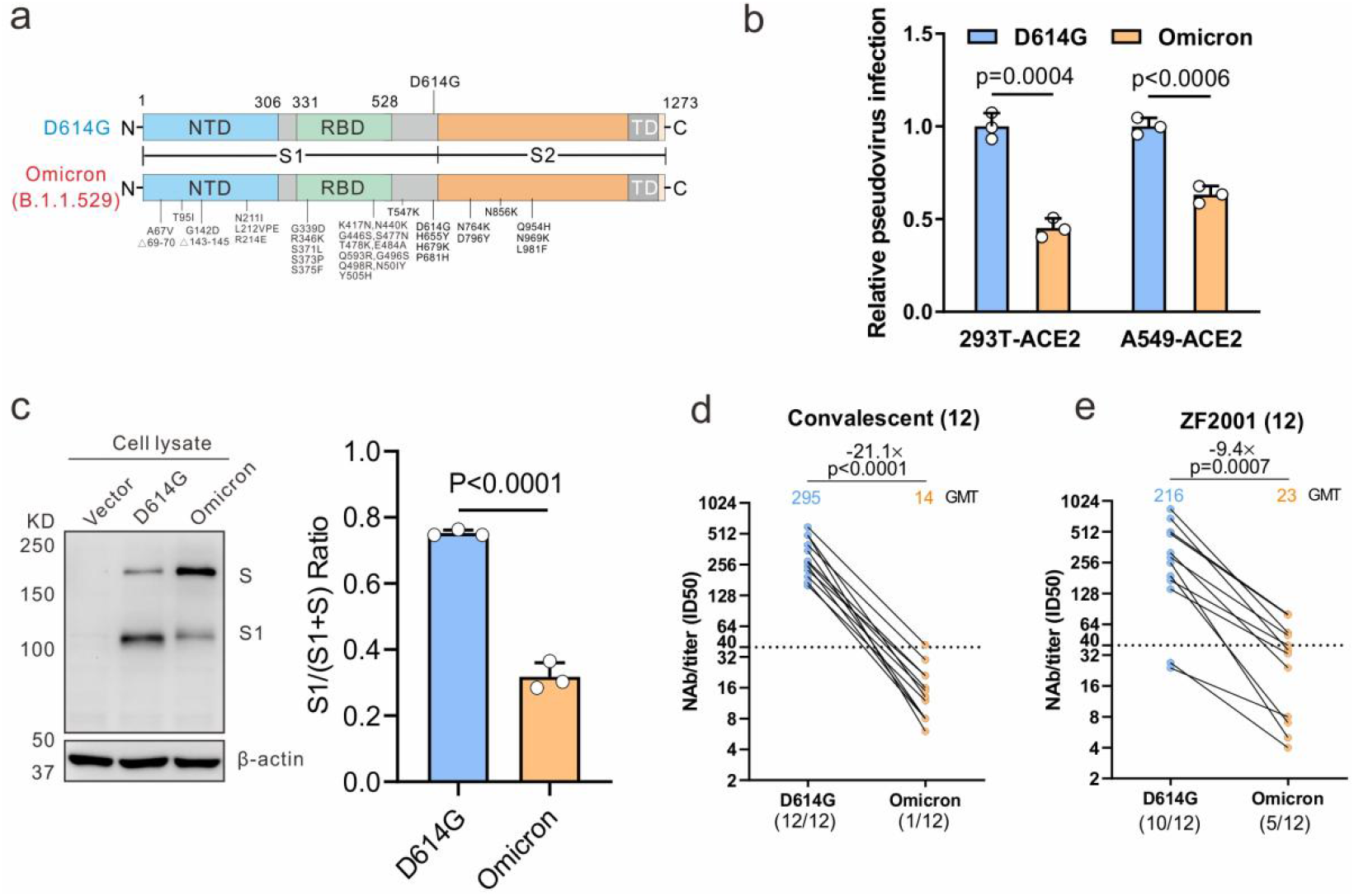
The effect of the Omicron variant on viral infectivity and immune escape. **a** Mutational landscapes in variants Omicron and D614G. **b** HEK293T-hACE2 cells or A549-hACE2 cells were infected with pseudotyped Omicron and D614G particles bearing all S mutations to determine viral infectivity using a luciferase assay. **c** HEK293T cells were transfected with the spike-expressing plasmids of Omicron and D614G variants and the immunoblots were probed with the indicated antibodies; the cleavage quantified by densitometry using ImageJ. **d-e** Pseudovirus-based neutralizing assays were performed to detect neutralizing activity of the sera from the convalescents (n=12, sampled at around 30 days after symptom onset) (**d**) and vaccinated individuals with RBD subunit vaccine ZF2001 (n=12, sampled at 15-60 days post the 3rd dose) (**e**) against SARS-CoV-2 Omicron and D614G variants. The half-maximal inhibitory dose (ID_50_) was indicated by geometric mean titers (GMT). The threshold of ID_50_ detection was 1:40. Statistical data analysis was performed using GraphPad Prism version 8.0 software. Student’s *t*-tests were used to compare between Omicron and D614G variants. Statistical significance was determined using ANOVA for multiple comparisons. When *P* values < 0.05, differences were deemed as the statistically significant.

## Results

To determine the effect of Omicron S protein mutations on its capacity of infectivity and immune evasion, a previously described method was used to construct Omicron pseudotyped virus^5^. After codon optimization according to the sequence from GISAID (EPI_ISL_6640917), the synthesized S gene of Omicron was constructed into a luciferase-expressing pseudotyping lentiviral system based on HIV-1 backbone. Pseudotyped Omicron particles bearing all S mutations were used to infect the HEK293T cells or A549 cells expressing ACE2 receptor for the measurement of viral infectivity in a luciferase assay. A previously constructed SARS-CoV-2 pseudovirus with the ancestral D614G mutation in S protein was also applied as the reference, since the D614G mutation has been reported among almost all SARS-CoV-2 variants. As shown in Fig. 1b, the lower entry efficiency of Omicron than D614G was observed with 49% and 37% decreased luciferase activity in HEK293T-hACE2 and A549-hACE2 cells at 72h post infection, suggesting that Omicron S protein mutations lead to its reduced infectivity. Furthermore, we transfected HEK293T cells with the S-expressing plasmids of Omicron and D614G variants to examine protein expression and cleavage. The immunoblot analysis displayed S proteins cleaved into two major bands denoting full-length S and S1 subunit by the cellular proteases. Compared with the expression level of the two proteins in the D614G variant, Omicron showed a reduced level of S1 subunit (Fig. 1c). It indicates less proteases cleavage of the Omicron variant, consistent with another report^6^.

On the other hand, pseudotype-based neutralizing assay was performed as previously described to analyze neutralizing antibodies (NAbs) elicited by previously infection or the RBD-based protein subunit vaccine ZF2001 against the Omicron variant^7^. Sera sampled at around one month after symptom onset from 12 convalescents who were previously infected by SARS-CoV-2 original strain shows a more than 20-fold decrease of neutralizing activity against Omicron variant, when compared to D614G variant (Fig. 1d). Only one of these 12 individuals remains slight neutralizing effect on the Omicron variant. Among 12 individuals vaccinated by RBD-based protein subunit vaccine ZF2001, 58.3% (7/12) sera sampled at 15-60 days after 3rd-dose vaccination did not neutralize Omicron. Geometric mean titers (GMTs, 50% inhibitory dose [ID_50_]) of these sera against Omicron were 9.4-fold lower than against D614G (Fig. 1e).

## Discussion

Here we have shown that the new SARS-CoV-2 variant Omicron S protein with a large number of mutations has an outstanding effect on the viral infectivity and immune escape ability. Unexpectedly, different with other VOCs, reduced entry efficiency and less cleavage ability were observed in our study. Further investigation using authentic virus should be addressed to validate and explain about Omicron’s infectivity and fast transmission.

Consistent with other studies, we have observed that protective immunity after previous infection could barely neutralize Omicron. Even worse, almost all vaccines which have extensively used exhibit the remarkable reduced neutralization against Omicron^8, 9^. However, it is worth noting that administration of a booster dose and vaccination of individuals with previous infection present a better neutralizing response^9^. In our study, the better neutralization of third-dose RBD subunit vaccine sera against Omicron displays a lower fold-change (9.4 folds) than convalescent sera (21.1 folds). We hypothesize that antibody affinity maturation induced by vaccines with multiple doses will be benefit for increased neutralization against future variants like Omicron^10^.

Taken together the results suggested a higher risk of breakthrough infections and reduced efficiency of the protective immunity elicited by existing vaccines. There are important implications about the modification and optimization of the current epidemic prevention and control including vaccine strategies and therapeutic antibodies against the new SARS-CoV-2 variant Omicron.

## Acknowledgements

We acknowledge funding support from the China National Natural Science Foundation (grant no. U20A20392), the 111 Project (No. D20028), the Research Fund Program of the Key Laboratory of Molecular Biology for Infectious Diseases, CQMU (No. 202105), the Emergency Project from the Science & Technology Commission of Chongqing (cstc2020jscx-fyzx0053), the Emergency Project for Novel Coronavirus Pneumonia from the Chongqing Medical University (CQMUNCP0302), the Leading Talent Program of CQ CSTC (CSTCCXLJRC201719), and a Major National Science & Technology Program grant (2017ZX10202203) from the Science & Technology Commission of China, National Natural Science Foundation of China (Grant No.82102361), China Postdoctoral Science Foundation (2021M693924), Natural Science Foundation of Chongqing, China (cstc2021jcyj-bshX0115) and Chongqing Postdoctoral Science Special Foundation (2010010005216630).

## Author Contributions

A.H., N.T., K.Wang., P.P. and J.H. developed the conceptual ideas and designed the study. J.H., P.P. and K.Wu performed the experiments and statistical analysis. Q.L. and J.C. provided the samples. All authors provided scientific expertise and the interpretation of data for the work. P.P. drafted the manuscript. All authors contributed to critical revision of the manuscript for important intellectual content. All authors reviewed and approved the final version of the report.

## Conflict of Interest

The authors declare no conflicts of interest.

## Notes

### Competing Interest Statement

The authors have declared no competing interest.

